# Freezing seawater samples alters the community composition and richness inferred from eDNA metabarcoding

**DOI:** 10.64898/2026.11.13.764904

**Authors:** Rose-Lynne Savage, Rute Clemente-Carvalho, Evan Morien, Matthew A Lemay

## Abstract

Environmental DNA (eDNA) has become a common approach for surveying biodiversity, yet processing aquatic samples can sometimes be challenging in the field. When eDNA water sample processing is delayed during remote field collections, whole water samples can be stored frozen to minimize further eDNA degradation. However, the freezing and thawing processes may influence eDNA integrity thus affecting species identification. We split bulk seawater samples to compare the marine taxa detected from water that had been filtered immediately to water that had been stored frozen for one month. Using DNA metabarcoding targeting a 313bp section of the cytochrome c oxidase subunit I (COI) mitochondrial barcode gene, we evaluated the effect of freezing on the inferred marine invertebrate communities. We found that previously frozen samples had significantly lower species richness and different community composition compared to the non-frozen samples. Particularly, amplified sequence variants (ASVs) of the copepod *Pseudocalanus* were significantly more abundant in non-frozen samples. These data suggest that freezing water samples intended for eDNA analyses may impede species detection for select organisms and significantly alter the inferred community structure. Studies using pre-frozen water samples should consider downstream implications when analyzing eDNA biodiversity.

## INTRODUCTION

Over the past decade biodiversity monitoring has increasingly incorporated the use of environmental genetic material for species detection and identification. Approaches using environmental DNA (eDNA) have been developed to monitor the biodiversity of plants and animals by using DNA they shed or excrete into the surrounding soil, water, or air (Deiner et al., 2017; Ficetola et al., 2008; Taberlet et al., 2012). The implementation of eDNA-based tools for species detection has become a powerful method to survey biodiversity and to detect rare and invasive species (Gold et al., 2021; Holman et al., 2019). Molecular techniques have improved on traditional monitoring methods as environmental DNA samples are non-destructive to organisms and the environment, are relatively simple to collect, and require no taxonomic expertise.

There are continued efforts to optimize and standardize the best eDNA practices in environmental monitoring as sample collection, laboratory processing, and analytical methods can vary significantly between studies. In aquatic environments eDNA is extracted from water samples and specific molecular markers are targetted for species identification. The recovery and detection of eDNA may be influenced by multiple steps during sample processing, including water sample storage, DNA collection (e.g. by filtering, precipitation, or centrifugation), and DNA extraction methods (Deiner et al., 2015; Takahara et al., 2020). It is especially important to consider eDNA degradation after collection as multiple external factors including temperature, pH, UV exposure, and microbial activity influence eDNA decay rates (Strickler et al., 2015). Filtering water samples is one of the optimal practices to capture eDNA (Deiner et al., 2015; Spens et al., 2017) and ideally samples are filtered immediately or within 24 hours after collection to minimize eDNA degradation (Hinlo et al., 2017; Pilliod et al., 2013). Unfortunately, practitioners are often faced with the inability to filter water immediately during surveys due to obstacles such as high frequency sampling, unavailability of sterile space or materials, limited infrastructure for sample processing, or limited workforce available. Due to these situations, it is common for water samples to be stored for a period of time before being processed.

The degradation of eDNA can be minimized during the storage of whole water samples by keeping water samples refrigerated, frozen, or stored with a preservative (Jo et al., 2021; Pilliod et al., 2013; Takahara et al., 2013; Williams et al., 2016). The preferred storage method will depend on facility capacities, the reagents required, and how long until processing is possible. Freezing whole water samples is a favorable option as it is simple, only requires freezing capacity, and does not require additional preserving reagents. Although freezing water samples may help to minimize the eDNA degradation during longer storage periods, it is not fully understood how the freezing and thawing processes affect eDNA analyses and conclusions.

Regardless of the potential effects on species detection, freezing water samples can be used when samples need to be stored for long periods of time (e.g. to facilitate sample shipping, delayed processing in a clean lab). Freshwater samples stored frozen for up to 21 days had the most stable fish eDNA concentrations compared to replicate samples kept for the same time at room temperature or refrigerated, without the use of a preservative (Takahara et al., 2020). Hinlo et al. (2017) compared the effects of various water sample preservation and storage methods on freshwater fish eDNA detection and concluded that when samples cannot be filtered within five days whole water samples should be stored frozen to minimize eDNA decay. However, despite keeping eDNA concentrations stable for storage, freezing water samples may still reduce species detection. Takahara et al. (2015) suggested that the freeze-thaw processes may retain more PCR inhibiting materials in the extracted DNA, thus affecting species-specific detection.

Previous investigations of how freezing water samples may affect eDNA species detection have mainly been from freshwater environments, estuaries, or aquaria (Hinlo et al., 2017; Takahara et al., 2015; Turba et al., 2023), focused only on targeted species detection (Hinlo et al., 2017; Takahara et al., 2020, 2015), or to validate the use of added preservatives (Takahara et al., 2020). Here we investigate how freezing marine water samples for delayed sample processing will influence the eukaryotic community composition resolved from eDNA metabarcoding. Our goal was to evaluate how freezing and thawing seawater samples prior to filtering would affect the inferred diversity of marine invertebrate communities from coastal British Columbia, Canada. We split whole water samples in half to capture and compare eDNA that had been filtered immediately to the fraction of the same sample that had been stored frozen in bottles for one month prior to being thawed and filtered.

## MATERIALS AND METHODS

### Sample Collection

Sampling took place at the Hakai Institute Oceanographic Station QU38 (50.117 °S, 125.223 °W) on March 2, 2022. Twenty replicate 2 L surface water samples were collected in bottles that had been sterilized using bleach and UV light. The samples were stored in coolers with ice packs, transported back to the lab, and processed within 3 hours of collection. Each 2 L sample was inverted to mix and homogenize the seawater. One liter of seawater was immediately filtered onto a 0.22 µm Sterivex filter (Millipore), which were preserved with Longmire’s solution (100 mM Tris, 100 mM EDTA, 10 mM NaCl, 0.5 % SDS) at-20°C (Longmire et al., 1997; Renshaw et al., 2015). These samples are hereon referred to as the non-frozen samples. The remaining 1 L fractions of seawater from each sample were transferred to sterile 1.5 L bottles and kept frozen at-20°C for 30 days. To minimize DNA decay by UV exposure and warm temperature during thawing, the samples were kept in coolers at room temperature for 48 hours to thaw. One liter of the thawed seawater samples was inverted, filtered onto a 0.22 µm Sterivex filter, and preserved with Longmire’s solution at-20°C. These samples will be referred to as the frozen samples.

This resulted in twenty paired replicates of non-frozen and frozen samples. The preserved Sterivexes from each non-frozen and frozen treatment were stored at-20°C for one day before DNA extraction.

### DNA Extraction

Extractions were performed in clean rooms designated for isolating DNA from environmental samples. An unused sterile Sterivex filter was included in the DNA extraction procedure as a negative control when processing each batch of samples. The DNA extraction procedure was based on a modified protocol using the Qiagen DNeasy Blood & Tissue extraction kit with a bead beating step for Sterivex filters (Kellogg et al., 2024a; Thomsen et al., 2012). The Sterivexes were thawed and the outlet placed in a 2 mL collection tube. Both the Sterivex and the 2mL collection tube were placed in a 50 mL conical tube and centrifuged at 2500 g for 1 minute to remove the Longmire’s solution from the Sterivexes. Approximately 0.3 g of 0.4 mm silica beads and 720 µL of ATL buffer were added to the Sterivexes. The Sterivexes were vortexed at maximum speed using a horizontal adaptor for ten minutes and then incubated at 56°C on rotation for thirty minutes. The bead beating and incubation steps were repeated a second time to ensure complete cell and particle disruption. Next, 80 µL of proteinase K was added to the Sterivexes and incubated on rotation at 56 °C for two hours. The lysate was collected through the inlet of the Sterivexes at a speed of 4200 g for 1 minute using the centrifugation method previously described. The manufacturer’s protocol for the DNeasy Blood & Tissue extraction kit, including a RNase digestion step, was followed for the remaining extraction. DNA was eluted in 200 µL of AE buffer, quantified using a Qubit 2.0 fluorometer and the Qubit™ dsDNA HS Assay kit, and stored at-20°C until PCR library preparation. DNA samples from each treatment were run on a QIAxcel capillary electrophoresis device using the DNA High Resolution kit and the Large-Fragment Marker kit to visualize DNA quality of frozen and non-frozen samples.

### Library Preparation

DNA metabarcoding for all samples and negative controls was carried out to amplify a short section of the mitochondrial cytochrome oxidase subunit 1 (COI) gene (mICOIintF and dgHCO2198R) (Leray et al., 2013; Meyer, 2003). All genomic processing was performed in a designated clean room for the preparation of eDNA libraries and was carried out using a two-step PCR method (Bourlat et al., 2016; Kellogg et al., 2024b).

The PCRs were conducted in 25 µL reactions including 12.5 µL of 2X Taq FroggaMix, 2 µL of 10 mg/mL BSA, 0.6 uL of 10uM forward and reverse primers containing Illumina overhangs, 2 µL of non-diluted DNA template, and nuclease-free water. The thermocycler conditions began with a 5 minute denaturation at 95°C followed by 40 cycles of 95°C denaturation for 30 seconds, primer annealing at 50°C for 30 seconds, and elongation at 72°C for 45 seconds. These reaction cycles were concluded with a final elongation at 72°C for 5 minutes. For each sample, triplicate PCRs were amplified and then pooled together and the pooled products were purified using SPRI magnetic beads (GE Healthcare) following the manufacturer’s protocol. The second PCR used the purified amplicon product from the gene specific PCR as a template and ligated the remaining Illumina sequencing adaptors and sample specific indexes required for dual-indexed metabarcoding. Indexing PCRs, reactions were conducted in 25 µL and included 12.5 µL of 2X Taq FroggaMix, 2.5 µL of 10 µM forward and reverse primers with adaptors and indexes, 2.5 µL of PCR product from the first PCR, and nuclease-free water to make up a 25 µL reaction mixture. The initial denaturation at 95°C ran for 3 minutes followed by 8 cycles of 30 second denaturation at 95°C, 30 seconds of primer annealing at 55°C, and 30 seconds of elongation at 72°C. A final 5 minute elongation at 72°C concluded the PCR. The indexed amplicon products were purified using SPRI magnetic beads and quantified using the Quant-iT™ dsDNA Assay Kit. Each sample was pooled at equimolar values. The resulting pooled library was sequenced using Illumina MiSeq Reagent kit V2 (500 cycles). We loaded 5 pM library spiked with 10% PhiX control (Illumina).

### Bioinformatics

We processed paired-end metabarcoding reads using the dada2 library v1.22.0 in R v4.1.3 (R Core Team, 2021), and used cutadapt v2.10 to trim primers (Martin, 2011). We filtered primer-trimmed reads using default dada2 parameters, except for these differences: we set trimRight to (25,25), truncLen to (0,0), minLen to (110,110), and maxEE to (6,8). After learning error rates, dereplication, sample inference, and read merging with dada2 using default parameters, we removed samples with fewer than 50 amplicon sequence variants (ASVs) present, and ASVs present in only a single sample with lower than 0.001 relative abundance (hereafter referred to as ‘singleton ASVs’). We removed bimeras (chimeric sequences with two identifiable sources) using default dada2 parameters. We used NCBI’s blastn 2.10.0+ to derive taxonomy assignments. COI sequences were blasted against two databases: NCBI NT database, plus a custom filtered set of COI sequences drawn from NCBI and BOLD (Ratnasingham and Hebert, 2007). We then combined the blast outputs. Blast output was limited to hits that had ≥ 96% similarity to the query, ≥ 50% query coverage, and an e-value of ≥ 1e-5, all other parameters were set to defaults. We used Galaxy Tool LCA (https://github.com/naturalis/galaxy-tool-lca) to determine LCA (last common ancestor) taxonomy strings for each ASV. The top scoring blast hit was applied without using LCA if it had ≥ 98% similarity with the query, otherwise, LCA was employed to determine a consensus taxonomy.

We used taxonomic filters to remove all ASVs that lacked a phylum-level identification as well as non-eukaryote ASVs, ASVs from vertebrates, and any putatively terrestrial ASVs (e.g. insects). The COI data was then split into two datasets for statistical analysis; the first consisted of all marine taxa identified in the sample (including marine invertebrates and algae), the second was a refined set of only ASVs that annotated to marine invertebrate taxa.

### Statistical analysis

All statistical analyses and graph generation were conducted in R version 4.1.2 (R Core Team, 2021). For each sample the Chao1 diversity index (Chao, 1984) was calculated to compare the richness of ASVs between treatments. An Analysis of Variance (ANOVA), including test with the paired sample replicates included as a random factor, was used to determine if ASV richness was significantly different between the non-frozen and frozen pre-processing treatments. The assumption of normality was validated using a Shapiro-Wilk normality test and homogenous variance was confirmed using the Levene test. A non-parametric Kruskal-Wallis rank sum test was used to determine if the average DNA concentration from non-frozen and frozen samples were also significantly different. Test results for assumption of normality of residuals and variance homogeneity for ANOVA are reported in the supplementary data (Table S1).

Statistical testing for differences in community composition and variance between pre-processing treatments were conducted using the vegan package version 2.6-4 (Oksanen et al., 2026). The *adonis2* function was used to determine if ASV communities were significantly different between treatments using the samples’ bray-curtis dissimilarity distances. The *betadisper* function was also used to determine if the samples from each treatment had different variances, potentially influencing PERMANOVA results.

Differential abundance analysis was used to identify ASVs that were significantly more abundant in either the frozen or non-frozen treatments. The phyloseq v1.38.0 R package (McMurdie and Holmes, 2013) and Wald t-tests with the DESeq2 package (Love et al., 2014) were used to identify differentially abundant ASVs between sample treatments.

## RESULTS

Sequencing of COI libraries produced 9,039,789 reads, of which 7,406,554 were retained after bioinformatic processing for subsequent filtering and analyses, with an average coverage of 184,922 reads per experimental sample. Taxonomic assignment identified 870 classified ASVs of marine eukaryotic taxa, of which 717 ASVs were classified marine invertebrates. Rarefaction curves show that sequencing effort captured the ASV diversity of all samples (Fig. 1).

**Figure 1.**
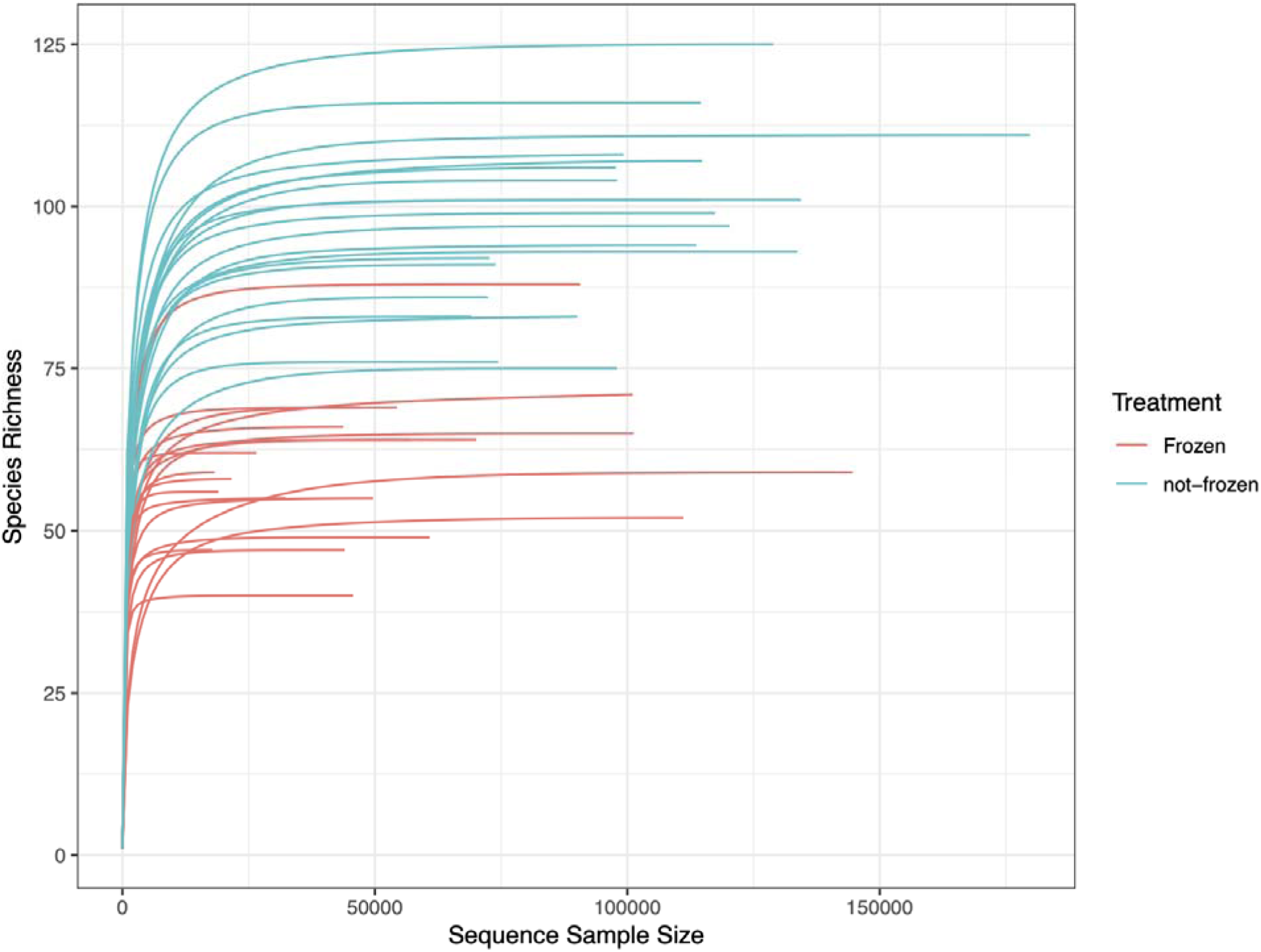
Rarefaction curves of the COI amplified sequence variants (ASVs) sequenced from non-frozen and frozen samples. Non-frozen sample curves are blue curves and orange curves indicate frozen samples.

### Relationship between DNA concentration and quality

Of the unclassified reads that were removed from further analysis because they lacked phylum-level annotation, there were more unassigned ASV reads in water samples that had been frozen relative to those which were not frozen. The average concentration of extracted DNA was significantly lower (p = 2.049 × 10^-7^, chi-squared = 26.986, df = 1) and relatively less variable in non-frozen samples, 1.1 ± 0.2 ng/µL, compared to the frozen samples, 2.7 ± 2.1 ng/µL. Higher DNA concentration from frozen samples did not result in greater ASV diversity. QIAxcel capillary electrophoresis showed that both frozen and non-frozen samples had degraded DNA. The non-frozen samples had greater consistency in DNA detection and fragment size across samples, while the frozen samples showed a broad variation in DNA degradation and in multiple samples DNA was not detected (Fig. S1).

### ASV Richness

For all the marine eukaryotes identified in the COI library, 180 ASVs (21% of ASVs, 56% of genera) were identified in both the frozen and the non-frozen treatments whereas 491 (56%) and 199 (23%) ASVs were unique to the non-frozen and frozen treatments, respectively (Fig. 2A). ANOVA detected significantly higher ASV richness in the non-frozen samples compared to the frozen sample treatment (Fig. 3A, p = 1.032 × 10^-10^, F = 160.48, NumDF = 1, DenDf = 19). This difference was also reflected in the ASV rarefaction curves where non-frozen samples appeared to capture a higher diversity of ASVs overall. Similar to the full dataset of all marine taxa, for the subset of marine invertebrates, the majority of ASVs were identified in non-frozen samples (Fig. 2B). There was also significantly higher ASV richness of marine invertebrates observed from the non-frozen treatment (Fig. 3B, p = 1.281 × 10^-6^, F = 48.23, NumDF = 1, DenDf = 19).

**Figure 2.**
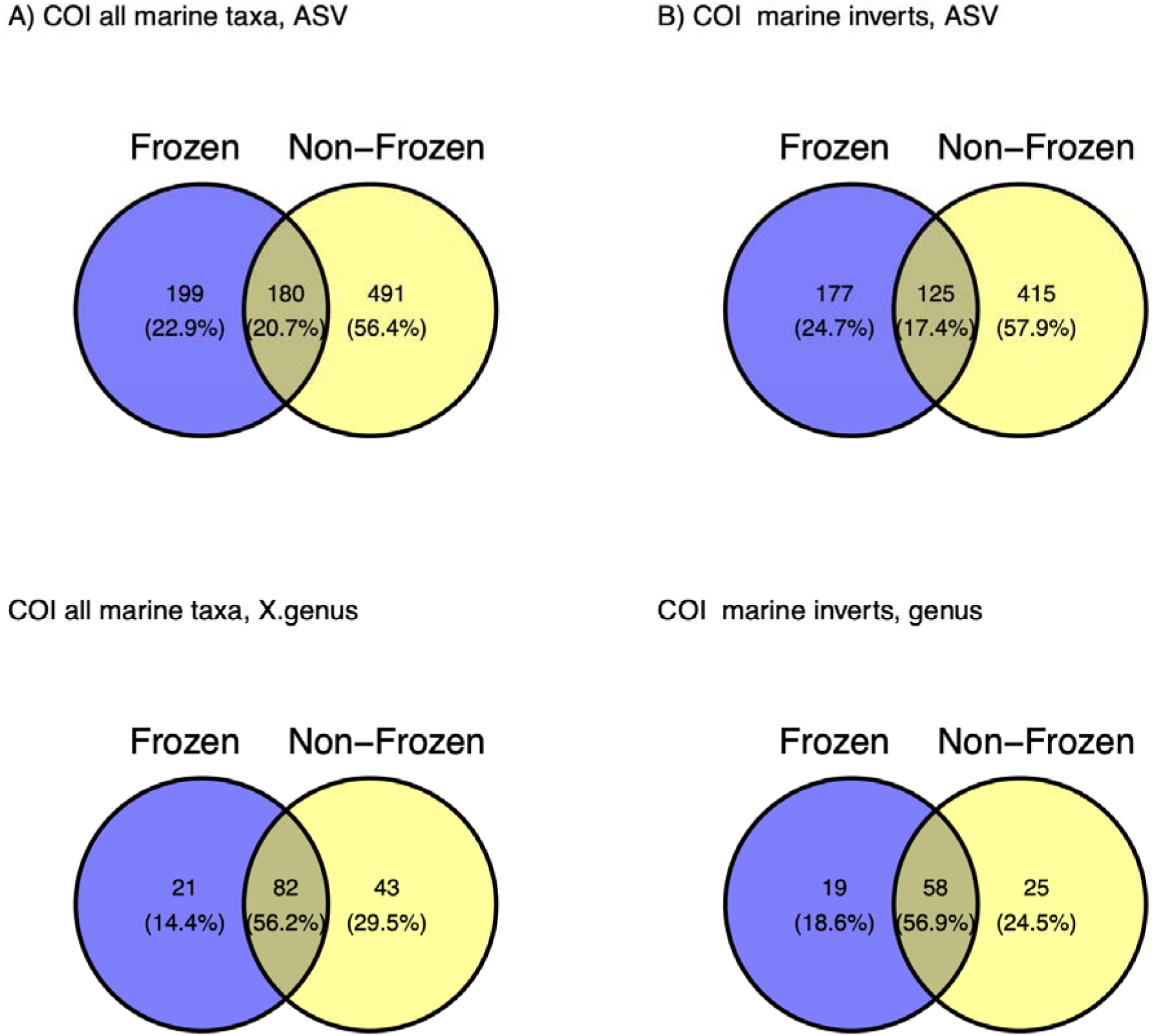
Venn diagrams of amplified sequence variants (ASVs, top) and classified genera (bottom) reads from non-frozen and frozen samples. Diagrams shown for cytochrome c oxidase subunit I (COI) reads of all marine taxa (A), COI reads of only marine invertebrates (B).

**Figure 3.**
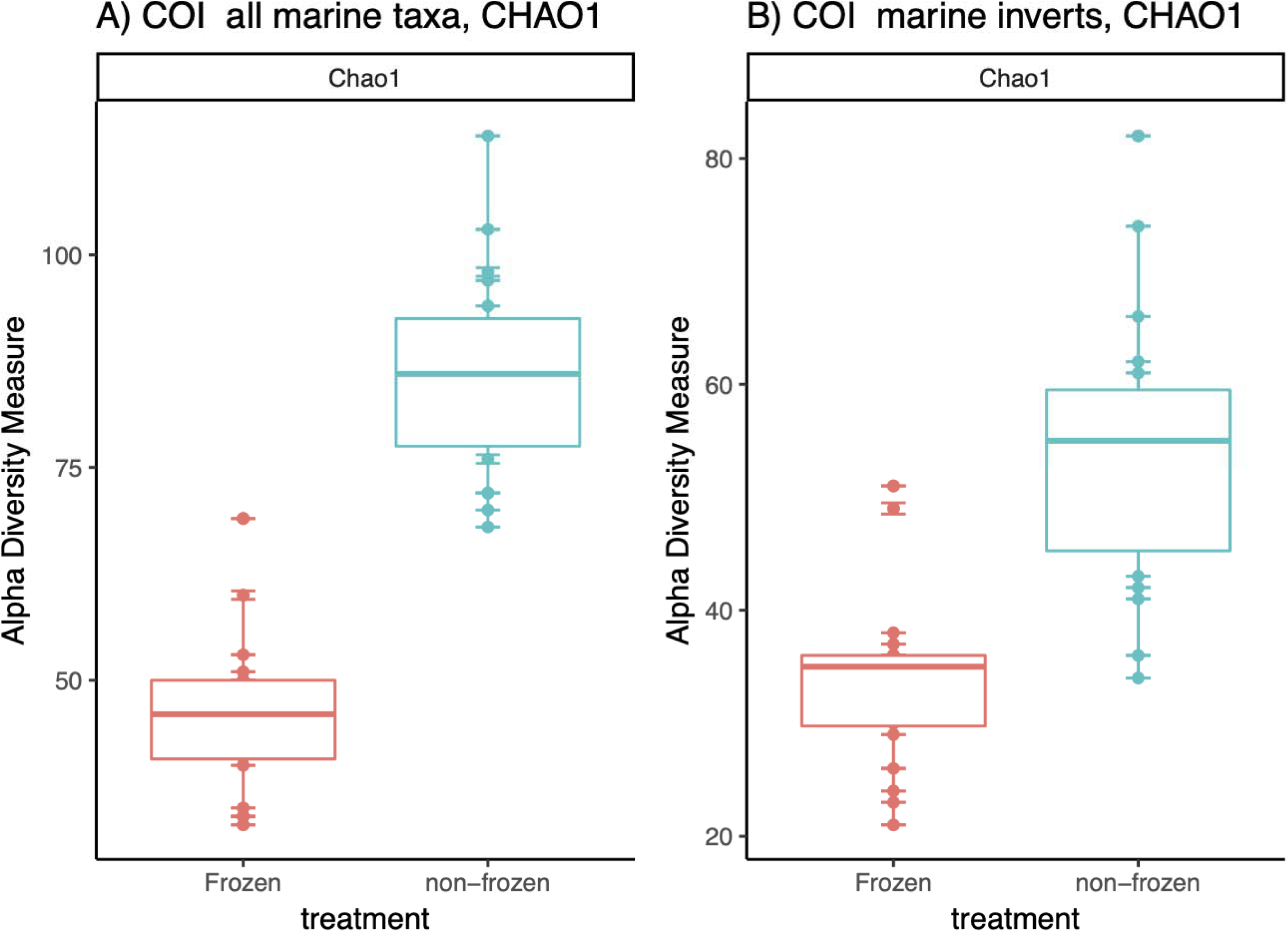
Chao1 diversity index of amplified sequence variants (ASVs) from cytochrome c oxidase subunit I (COI) ASVs of all marine taxa (A), COI ASVs of only marine invertebrates (B). The mean ASV richness is indicated by the centerline in the boxplots. The asterisks indicated a significant difference in Chao1 diversity between treatments.

### Community Composition and differential abundance

PERMANOVA indicated that COI communities were significantly different between the non-frozen and frozen treatments for all the marine taxa (*adonis2*, p = 0.001, F = 15.033, df = 39) and for only the marine invertebrates detected (*adonis2*, p = 0.001, F = 11.8, df = 39; Fig. 4A, B). However, the significant PERMANOVA could be influenced by the different sample variances of the treatments (*betadisper,* all marine taxa: p = 0.001, F = 112.91, df =38 and only invertebrates: p = 0.001, F = 81.72, df =38). Regardless of the differences in variances, pre-processing treatment is likely driving community differences due to the distinct separation of the treatment clusters.

**Figure 4.**
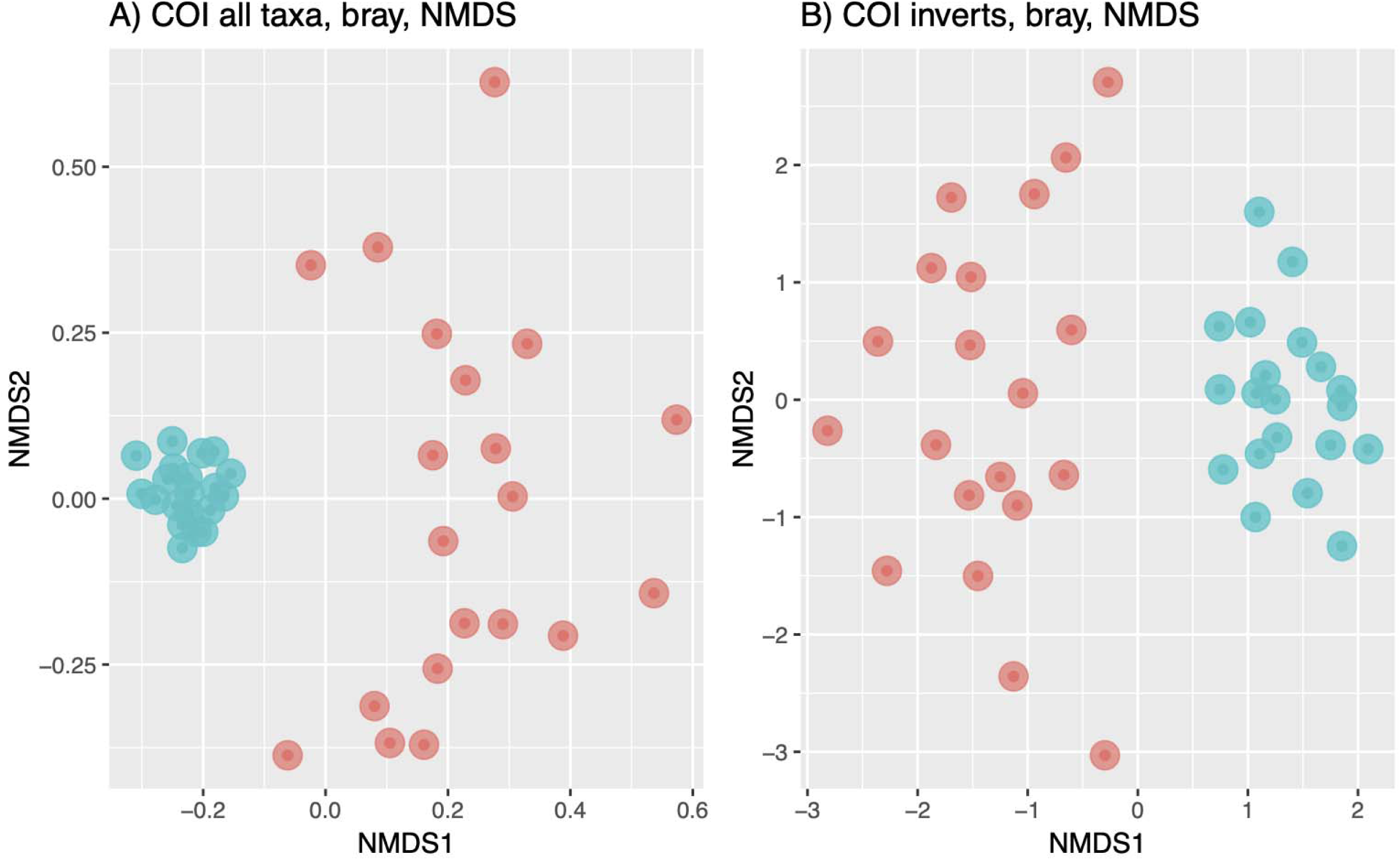
Non-Metric Multidimensional Scaling (NMDS) of the cytochrome c oxidase subunit I (COI) of all marine taxa (A), COI of only marine invertebrates (B) based on Bray-Curtis dissimilarity indexes. Non-frozen samples are blue circles and frozen samples are orange circles.

Marine diversity as inferred from COI in non-frozen samples was fairly consistent among replicates (Fig. 5). These non-frozen marine eukaryotic communities were mainly dominated by ASVs classified as the haptophyte *Phaeocystis* and the calanoid copepod *Pseudocalanus* (Fig. 5). The top ten ASVs with the highest relative abundance from the non-frozen samples belonged to the genera *Skeletonema* (Ochrophyta), *Pseudochattonella* (Ochrophyta), *Balanus* (Arthropoda), *Evasterias* (Echinodermata), and *Strongylocentrotus* (Echinodermata). In contrast, frozen samples were much more variable across replicates and were dominated by ASVs of similar genera including *Evasterias, Balanus*, *Phaeocystis*, and *Strongylocentrotus* (Fig. 5).

**Figure 5.**
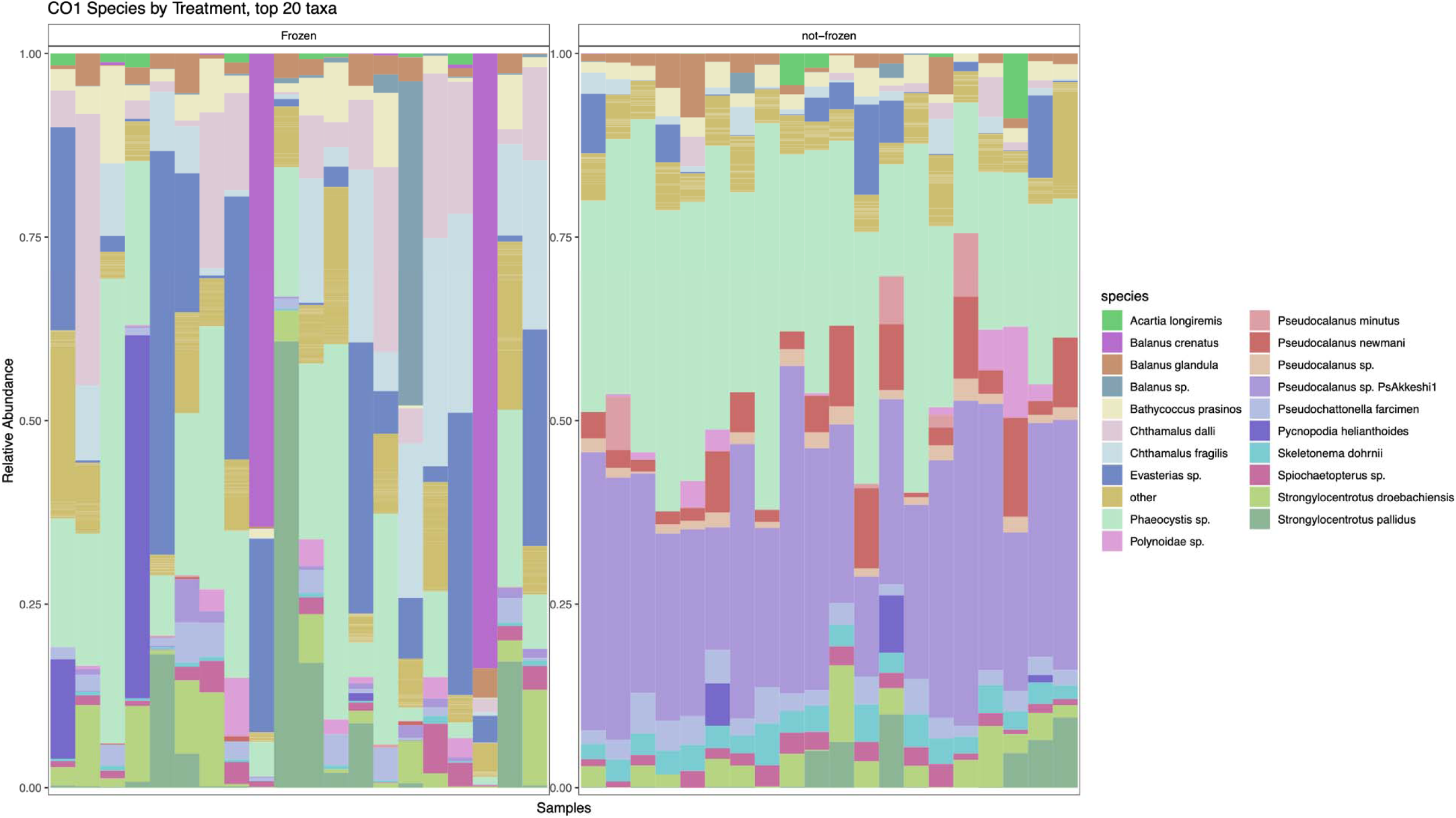
The relative abundance of the top twenty most abundant cytochrome c oxidase subunit I amplified sequence variants (ASVs) from samples of either non-frozen and frozen pre-processing treatment. Sequence variants are coloured by species.

Marine eukaryotic ASVs identified by differential abundance analysis as being increased in the non-frozen samples included *Pseudocalanus* (copepod), *Thalassionema* (diatom), *Pleurosigma* (diatom), *Thalassiosira* (diatom), *Skeletonema* (diatom), *Nitzschia* (diatom), *Prorocentrum* (dinoflagellate), *Heterocapsa* (dinoflagellate)*, Mesochaetopterus* (polychaete), *Dictyocha* (flagellate), *Pseudochattonella* (flagellate), *Leathesia* (algae), and *Phaeocystis* (algae) (Fig. 6, Fig.S2). Conversely, frozen samples had significantly higher abundances of an Asteriidae (sea star) ASV, *Balanus* (barnacle) and *Chthamalus* (barnacle), and the *Glycinde* (polychaete).

**Figure 6.**
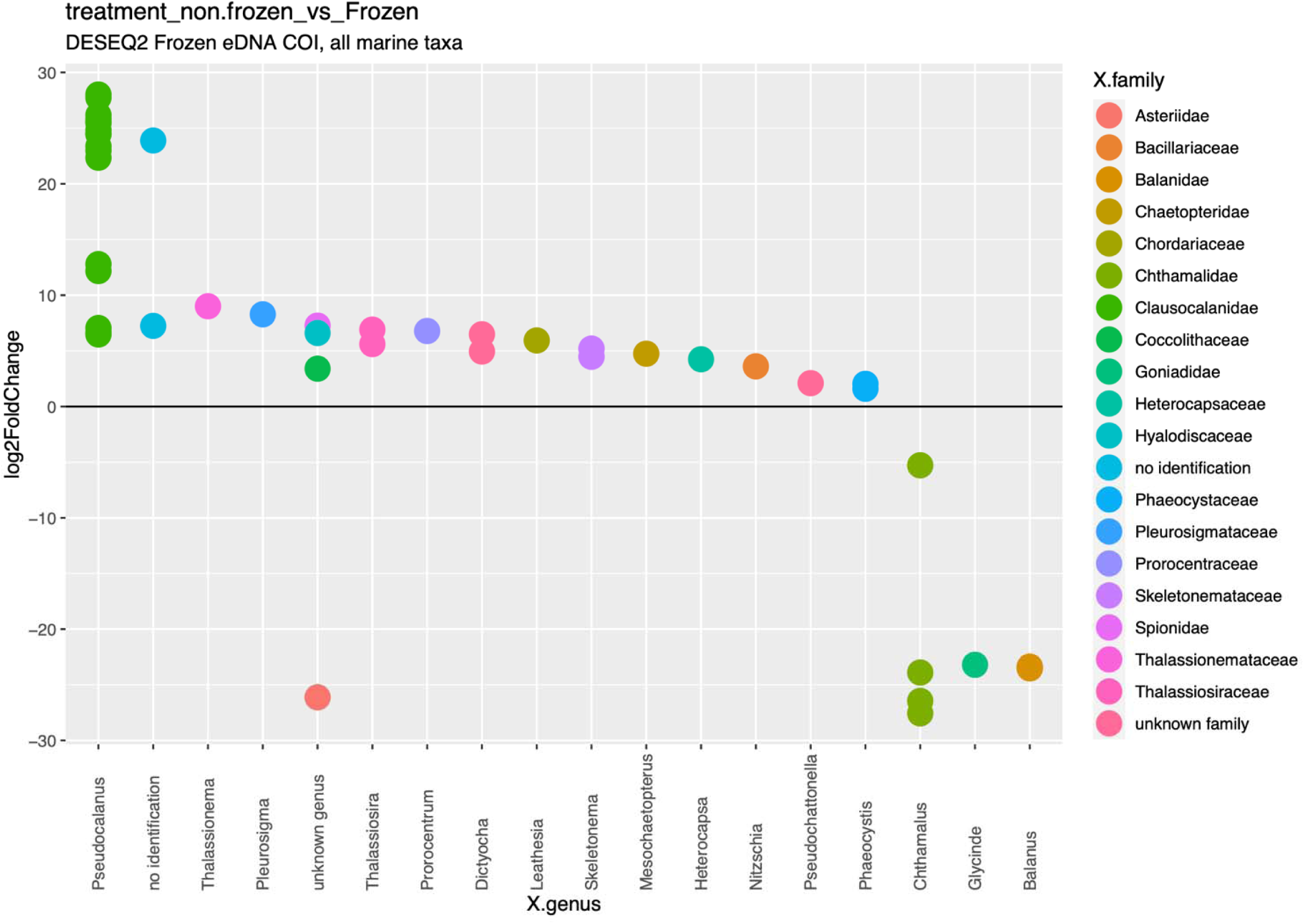
Differential abundance analysis using DESeq2 of amplified sequence variants (ASVs) from cytochrome c oxidase subunit I amplicon libraries. Each point represents the log fold change in abundance of differentially abundant ASVs identified in frozen or non-frozen samples. ASVs with a positive log fold change (above) are more abundant in non-frozen samples and samples with a negative log fold change (below) are more abundant in frozen samples.

## DISCUSSION

Immediate water filtering is not always feasible during field research and water samples are often frozen to facilitate shipment or delayed processing. Our results illustrate that for eDNA metabarcoding surveys, freezing seawater samples prior to processing can significantly alter the inferred biodiversity compared to samples that are immediately filtered. We observed significant reduction in diversity of marine eukaryotic communities resolved from cytochrome oxidase subunit 1 (COI) gene metabarcoding, as well as significant differences in community composition, with many taxa differentially represented between frozen and non-frozen samples. In addition, freezing bulk water samples introduced greater variation in DNA degradation, potentially increasing DNA decay. Despite significantly higher concentrations of extracted DNA from frozen samples, there was overall lower diversity relative to the paired non-frozen samples.

The number of classified marine COI ASVs was lower in water samples that had been stored frozen and the overall community composition was significantly different to that observed in non-frozen samples. *Pseudocalanus* copepod ASVs in particular were dominant in non-frozen samples and were detected at a much lower relative abundance in the frozen samples. Due to this potential sensitivity of *Pseudocalanus* eDNA to freezing, we recommend that water samples should not be frozen when aiming to target populations of this copepod and alternative preservation methods should be considered when water processing is delayed for longer than 24 hours (Takahara et al., 2020; Williams et al., 2016). The significant reduction of *Pseudocalanus* ASVs in frozen samples likely inflated the relative abundance of the remaining community ASVs, as is a caveat of the proportional nature of DNA metabarcoding data. This could be responsible for ASVs detected at a higher abundance in frozen samples by differential abundance analysis. Therefore, when working with samples that have been frozen and thawed it will be important to consider this potential influence on the overall diversity and on select taxa.

Freezing water samples has been shown to reduce qPCR detection of fish from freshwater eDNA samples, potentially due to PCR inhibitors retained in extracted DNA from the freezing and thawing processes (Takahara et al., 2015). PCR inhibitor retention may be responsible for the reduced COI diversity observed in frozen samples. Inhibitors can interfere with PCR in various ways (eg. by template DNA binding or Taq polymerase inhibition). Opel et al. (2010) demonstrated that PCR inhibition by DNA binding inhibitors can be influenced by short tandem repeat (STR) amplicon size as well as the primer melting temperature (T_m_). For example, primers with higher melting temperatures can bind more strongly to template DNA and outcompete inhibitor binding. DNA binding inhibitors also block primer elongation, which would make smaller amplicons less affected by these inhibitors (Opel et al., 2010). For amplicon library preparation we suggest testing previously frozen water samples for potential PCR inhibitors with methods to release PCR inhibition, including DNA clean up or extraction kits designed for inhibitor removal, DNA template dilution in PCR, addition of BSA (Deiner et al., 2015), or inhibitor resistant Taq polymerases (TaqMan Environmental Master Mix 2.0 (Life Technologies) used in (Takahara et al., 2015).

Many investigations have been carried out with bulk eDNA water samples to determine optimal long-term preservation strategies (Hinlo et al., 2017; Takahara et al., 2020; Williams et al., 2016). Aside from freezing, the other methods for long term water storage involve the addition of preserving buffers such as Longmires or benzalkonium chloride (Jo et al., 2021; Takahara et al., 2020; Williams et al., 2016). Freezing is an appropriate method for bulk water storage when sample processing must be delayed for longer than a couple days, however water sample properties may affect DNA degradation during frozen storage (Hinlo et al., 2017). Further work into longer storage periods (approximately greater than two months) should be conducted to determine if archiving bulk water samples can sustain environmental DNA yield and integrity.

We have demonstrated how freezing and thawing bulk water samples can influence species detection and community composition resolved by DNA metabarcoding. Reduced diversity should be considered when analyzing previously frozen eDNA water samples, especially for COI metabarcoding using these same primers. As eDNA methods develop and become incorporated with traditional monitoring methods, it will be important to determine how different steps in sample collection, processing, and library preparation bias can influence overall eDNA metabarcoding analyses.

## ACKNOWLEDGEMENTS

We would like to thank Kyle Hall and Tyrel Froese for assistance with field work and sample collection and the Tula Foundation for funding this work.

## DATA AVAILABILITY

Raw Illumina MiSeq data have been deposited at the NCBI Sequence Read Archive (SRA) BioProject ID: PRJNA937254. Figure S1, Figure S2, and Table S1 are available in the Supplemental Material.

## AUTHOR CONTRIBUTIONS

Rose-Lynne Savage (RLS) designed the experiment, collected the samples, processed and analyzed eDNA samples, and drafted the manuscript. Rute Clemente-Carvalho (RCC) contributed to the experimental design, assisted with lab processing, carried out Illumina sequencing. Evan Morien (EM) provided bioinformatics support and assisted with data analyses. Matthew A Lemay (MAL) contributed to experimental design and supervised the data collection and analysis. All authors contributed to writing the manuscript and have approved the final article.

