## Supplemental Material for "Freezing seawater samples alters the community composition and richness inferred from eDNA metabarcoding"

**Figure S1.** QIAxcel capillary electrophoresis of eleven non-frozen (A) and frozen (B) paired samples. The first well shows the Large Fragment Size Marker analyzed alongside the samples.

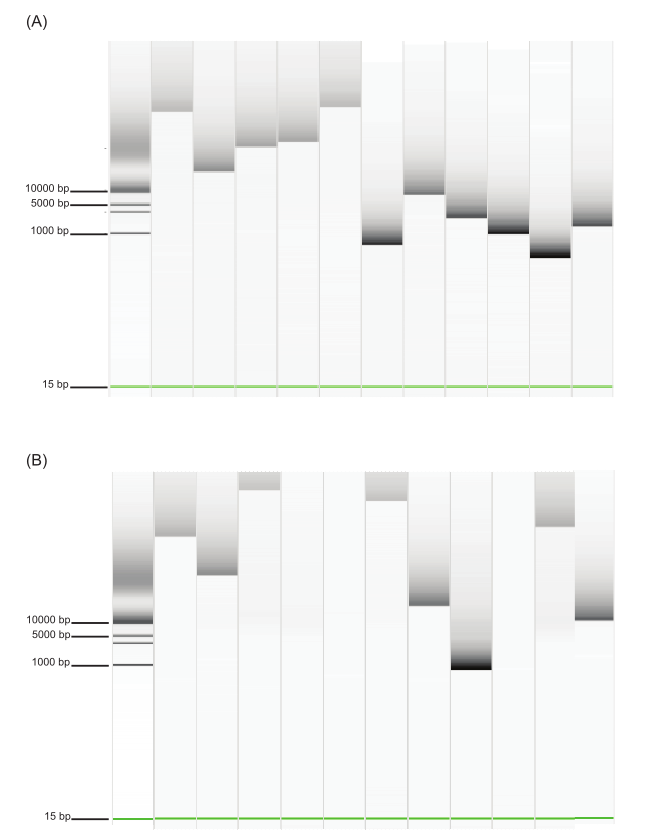

**Figure S2.** The relative abundance of the top twenty most abundant cytochrome c oxidase subunit I (COI) amplified sequence variants (ASVs) from samples of either frozen and non-frozen pre-processing treatment for each paired sample replicate.

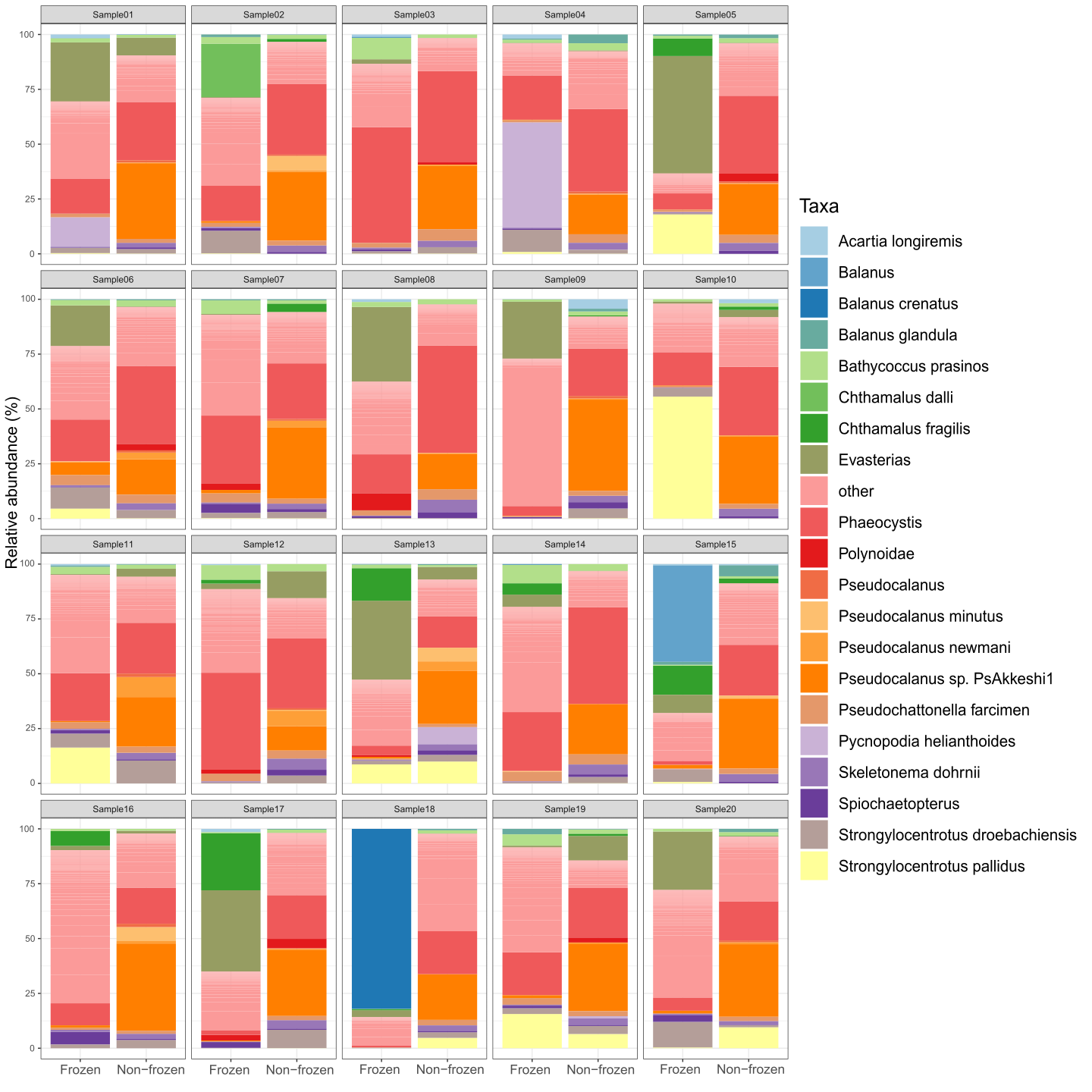

**Table S1.** Results for tests of assumptions of normality and homogeneous variance to continue with ANOVA tests.

|  | **ANOVA with sample replicate as random factor** | **Shapiro-Wilk test** | **Levene’s Test** | **Conclusion** |
| --- | --- | --- | --- | --- |
| **COI, all marine eukaryotic taxa** | F = 160.48  NumDf = 1  DenDF = 19  p = 1.032 × 10^-10^ | W = 0.97096  p = 0.3859 | F = 1.8392  Df = 38  p = 0.183 | Normality and equal variance assumptions met for ANOVA |
| **COI, marine invertebrates** | F = 48.23  NumDf = 1  DenDF = 19  p = 1.281 × 10^-6^ | W = 0.96772  p = 0.304 | F = 3.8311  Df = 38  p = 0.05769 | Normality assumption met for ANOVA. Equal variance met but is almost violated |
